# Analysis of SARS-CoV-2 RNA-Sequences by Interpretable Machine Learning Models

**DOI:** 10.1101/2020.05.15.097741

**Authors:** Marika Kaden, Katrin Sophie Bohnsack, Mirko Weber, Mateusz Kudła, Kaja Gutowska, Jacek Blazewicz, Thomas Villmann

## Abstract

We present an approach to investigate SARS-CoV-2 virus sequences based on alignment-free methods for RNA sequence comparison. In particular, we verify a given clustering result for the GISAID data set, which was obtained analyzing the molecular differences in coronavirus populations by phylogenetic trees. For this purpose, we use alignment-free dissimilarity measures for sequences and combine them with learning vector quantization classifiers for virus type discriminant analysis and classification. Those vector quantizers belong to the class of interpretable machine learning methods, which, on the one hand side provide additional knowledge about the classification decisions like discriminant feature correlations, and on the other hand can be equipped with a reject option. This option gives the model the property of self controlled evidence if applied to new data, i.e. the models refuses to make a classification decision, if the model evidence for the presented data is not given. After training such a classifier for the GISAID data set, we apply the obtained classifier model to another but unlabeled SARS-CoV-2 virus data set. On the one hand side, this allows us to assign new sequences to already known virus types and, on the other hand, the rejected sequences allow speculations about new virus types with respect to nucleotide base mutations in the viral sequences.

**Author summary:** The currently emerging global disease COVID-19 caused by novel SARS-CoV-2 viruses requires all scientific effort to investigate the development of the viral epidemy, the properties of the virus and its types. Investigations of the virus sequence are of special interest. Frequently, those are based on mathematical/statistical analysis. However, machine learning methods represent a promising alternative, if one focuses on interpretable models, i.e. those that do not act as *black-boxes.* Doing so, we apply variants of *Learning Vector Quantizers* to analyze the SARS-CoV-2 sequences. We encoded the sequences and compared them in their numerical representations to avoid the computationally costly comparison based on sequence alignments. Our resulting model is interpretable, robust, efficient, and has a self-controlling mechanism regarding the applicability to data. This framework was applied to two data sets concerning SARS-CoV-2. We were able to verify previously published virus type findings for one of the data sets by training our model to accurately identify the virus type of sequences. For sequences without virus type information (second data set), our trained model can predict them. Thereby, we observe a new scattered spreading of the sequences in the data space which probably is caused by mutations in the viral sequences.

## Introduction

The coronavirus disease 2019 (COVID-19) caused by SARS-CoV-2 viruses, whose origin lies probably in Wuhan (China), is a severe respiratory disease [1]. Currently (May 2020), it is spreading rapidly all over the world [2]. Yet there are several indicators that the molecular characteristic evolves during time [3, 4]. This evolution is mainly driven by mutations, which play an essential role and maybe accompanied by mechanisms of stabilization [5, 6].

The analysis of the genomic structure by sequencing is currently topic of ongoing research to better understand the molecular dynamics [7]. Obviously, changing the genomic structure may cause new properties and, hence, could increase the difficulties in finding drugs for treatment. For example, changes may lead to behavioral changes, such as the increased binding of the SARS-CoV-2 surface glycoprotein to human ACE2 receptors [8].

Viruses of the family *Coronaviridae* possess a single stranded, positive-sense RNA genome ranging from 26 to 32 kilobases in length and frequently are extremely similar [9]. Therefore, the analysis of those sequences to understand the genetic evolution in time and space is very difficult. This problem is magnified by incorrect or inaccurate sequencing [10]. Further, mutations are not equally distributed across the SARS-CoV-2 genome [11]. The molecular differences in coronavirus populations were investigated using phylogenetic trees so far resulting in three clusters which are identified as virus types [12]. Yet, SNP-based radial phylogeny-retrieved trees of SARS-CoV-2 genomes result in five major clades [11]. Generally, a disadvantage of those decision-tree-like approaches is the problem of out-of-sample considerations, i.e. new data cannot easily be integrated [13,14]. The respective tree has to be reconfigured completely, which frequently leads to major changes in the tree structure [15,16].

Frequent mutations in SARS-CoV-2 genomes are in the genes encoding the S-protein and RNA polymerase, RNA primase, and nucleoprotein. Applying a sequence alignment and similarity comparison using the Jaccard index, a method for monitoring and tracing SARS-CoV-2 mutations was established in [17]. However, a general mathematical evaluation of similarities is crucial because respective similarity measures only partially reflect all biological aspects of similarity between RNA sequences [18]. Alignment based methods usually rely on variants of the Levenshtein distance [19], which, however, are computationally costly: *O* (*n*_1_ · *n*_2_) is the time complexity for both the Needleman-Wunsch-algorithm [20] and for the Smith-Waterman-algorithm [21,22],where *n*_1_ and *n*_2_ are the sequences length. Hence, if *n*_1_ = *n*_2_ = *n* the complexity is simply *O* (*n*^2^). Both approaches solve internally a mathematical optimization problem, i.e. both algorithms belong to the algorithmic class of dynamic programming with high computational complexity. Other alignment based methods consider (multiple) longest common subsequences with similar complexity [23].

Therefore, alignment-free alternatives are promising [18,24–28]. Most common those approaches are *Bag-of-Words* (BoW, [29]), information theoretic methods based on the *Kolmogorov-Smirnov-complexity* [30] and the related *Normalized Compression Distance* [31,32]. Recently, similarities based on *Natural Vectors* gained attraction [33–35]. These methods have in common that the sequences are considered in terms of their statistical properties and distributions of the nucleotides. However, local information like precise nucleotide positions as well as specific motifs are lost. An overview of prominent measures and their behavior for sequence analysis can be found in [36,37].

In the present publication, we investigate whether alignment-free dissimilarities are suitable for the identification of SARS-CoV-2 clusters/classes in combination with *interpretable machine learning methods* for clustering and classification [38,39]. This we do for two data sets: GISAID-data and NCBI-data. For the first one, virus classes (types) were identified by phylogenetic tree analysis in [12], whereas the second one is without class information.

Although deep neural network approaches provide impressive results in sequence classification [16,40–42], deep architectures are at least difficult to interpret [43]. Therefore, we focus on applying prototype based methods using alignment-free dissimilarity measures for sequence comparison. In fact, prototype-based machine learning models for data classification and representation are known to be interpretable and robust [44–46]. Using such methods for the SARS-CoV-2 sequence data, first we verify the classification results for the GISAID-data. In particular, we classify the sequences by a learning vector quantizer, which is proven to be robust and interpretable [45, 47]. Thereafter, we use this model to classify the new data from the NCBI. Moreover, this interpretable classifier provides correlation information regarding data features contributing to a class discrimination. This additional knowledge allows a further characterization of the virus classes.

## Materials and methods

### SARS-CoV-2 Sequence Databases in Use

In order to investigate SARS-CoV-2 viruses in terms of sub-type spreading, two virus sequence data sets were considered.

### The GISAID Dataset *D_G_*

The first one, abbreviated by *D_G_*, is from the GISAID coronavirus repository (GISAID – Global Initiative on Sharing Avian Influenza Data). It consists by March 4th, 2020 of 254 coronavirus genomes, isolated from 244 humans, nine Chinese pangolins, and one bat *Rhinolophus affinis.* After preprocessing, 160 complete human sequences are obtained as described in [12], where these genomes of SARS-CoV-2 have been used to create a phylogenetic network. The resulting network analysis distinguished three types of the virus (cluster) *A, B,* and *C*: *A* is most similar to the bat virus, whereas *B* are sequences obtained from *A* by two mutations: the synonymous mutation T8782C and the non-synonymous mutation C28144T changing a leucine to a serine. A further non-synonymous mutation G26144T changing a glycine to a valine lead from *B* to type *C*. In this sense, the classes (virus types) code implicitly the evolution in time of the virus.

In our data analysis, we removed two sequences, whose accession numbers occur twice in the data record, and another two, which we identified as not human resulting in 156 final sequences. Additionally, we take the type/class information as label for the virus genome sequences and, hence, as reference. A detailed data description as well as complete list of sequences can be found in [12]. The virus type assignments and additional data (country, collection date) as well as accession numbers for all 156 sequences in use are additionally provided in the supplementary material.

The complete data information can be found in supplementary files S12 Data.

### The NCBI dataset *D_N_*

The second data set including 892 complete genomes has been selected from the National Center for Biotechnology Information (NCBI) Viral Genome database [48], and GenBank [49] by April 19th, 2020, see Tab. 1. These data are human based sequences and provide additionally the country information from which the sequences originate, as well as their collection date. For each sequence we have also derived a more general assignment to regions based on the country information, which includes the following values: USA, China, Europe, and Others. The accession number and the additional data used in the analysis have been included in the supplementary material. We refer to this data set by *D_N_*.

Remark, although the SARS-CoV-2 virus is an RNA virus, the sequences provided by databases are given using the DNA-coding. In the following, we take over this convention and do not explicitly refer to that later.

**Table 1.**
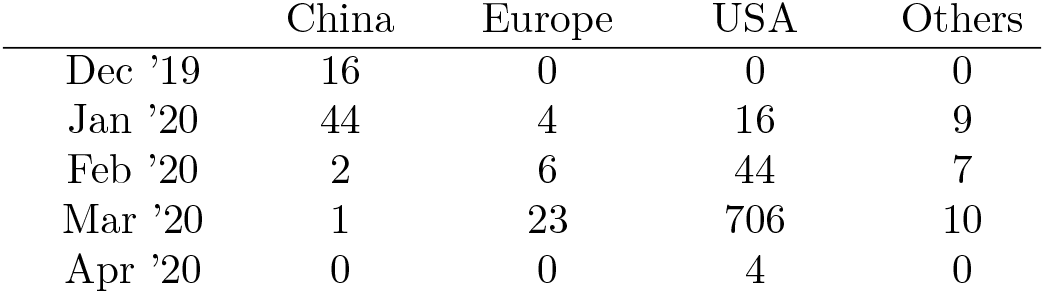
Distribution of the NCBI-data *D_N_* regarding regions and month of collection date.

|  | China | Europe | USA | Others |
| --- | --- | --- | --- | --- |
| Dec '19 | 16 | 0 | 0 | 0 |
| Jan '20 | 44 | 4 | 16 | 9 |
| Feb '20 | 2 | 6 | 44 | 7 |
| Mar '20 | 1 | 23 | 706 | 10 |
| Apr '20 | 0 | 0 | 4 | 0 |

Again, the complete data information can be found in supplementary files S12 Data.

### Representation of RNA Sequences for Alignment-free Data Analysis

Several approaches were published to represent sequences adequately for alignment-free comparison. These method range from chaos game representation to standard unary coding or matrix representations. An overview is given in [27] and [36, 37]. Here we focus only on two of the most promising approaches – *Natural Vectors* and *Bag-of-Words.*

### Natural Vectors

*Natural Vectors* (NV) for nucleotide sequence comparison is based a statistical sequence description for the distribution of nucleotide positions in a sequence **s** based on the alphabet 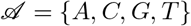 [33,34]. Let 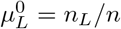 be the relative number (frequency) of the nucleotide 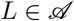 and *p_L_* (*j*) /*n, j* = 1…*n_L_* is the *relative* position of the *k*th nucleotide *L* in the sequence. Let *E* [*r*] further be the expectation operator of a random quantity *r*. Thus 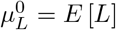. Further, we denote by 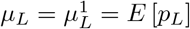 the mean relative position of the nucleotide *L* in the sequence. The *k*-th centralized moment 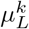 for *k* ≥ 2 is given as 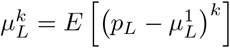. Then, the natural vector of order *K* for a sequence s is given as

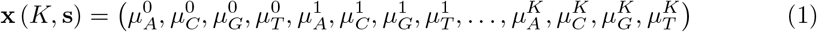

whereby we again drop the dependencies on *K* and s, if it is not misleading. Natural vectors are usually compared in terms of the *l_p_*-metric

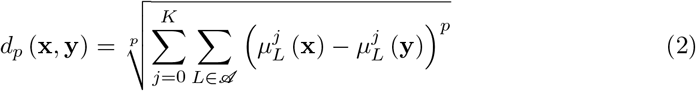

giving the Euclidean distance for *p* = 2. Kendall-statistics, as a kind of correlation measure, was applied in [50].

The NV-description of sequences can also be applied for nucleotide sequences containing ambiguous characters (degenerate bases) [35,51]. This yields an extended alphabet 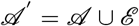. In that case, weights 0 ≤ *w_L_* (*s_i_*) ≤ 1 are introduced for each 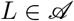 with

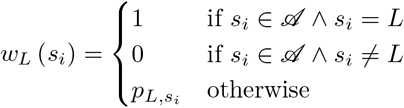

where *p_L,s_i__* is the probability that the detected ambiguous character 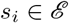 should be the character *L*. These weights have to be taken into account during the expectation value calculations [35].

### Bag-of-Words

Another popular method to compare RNA/DNA sequences is the method *Bag-of-words* (BoW) based on 3-mers, where the set *S* of words contains all possible 64 triplets defined by the nucleotide alphabet 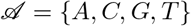 [24,25,27,41]. Thus all sequences s are coded as (normalized) histogram vectors of dimensionality *n* = 64, such that we have for each sequence the corresponding histogram vector **h** (s) ∈ ℝ^*n*^ with the constraints *h_k_* (s) ≥ 0 and 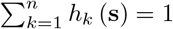. Mathematically speaking, these vectors are discrete representations of *probability densities*. If the latter constraint is dropped we have discrete representations of *positive functions.* The assignments of the triplets to the vector components *h_i_* is provided in the supplementary material. If it is not misleading we drop the dependence on s and simply write **h** instead of **h** (s). As for NV, nucleotide sequences with ambiguous characters can be handled using appropriate expectation values.

Obviously, comparison of those histogram vectors can be done using the usual Euclidean distance. However, motivated by the latter mentioned density property, an alternative choice is to compare by means of divergence measures [52]. In the investigations presented later, we applied the Kullback-Leibler-divergence [53]

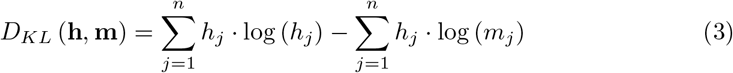

for sequence histograms **h** and **m**. Note that the first term in (3)) is the negative Shannon entropy 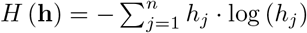 whereas 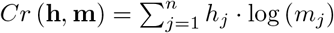 is the Shannon *cross-entropy.* Yet, other divergences like Rényi-divergences could be used [54]. We refer to [55] for a general overview regarding divergences in the context of machine learning.

The assignment of the nucleotide triplets to the histogram dimension can be found in the supplementary material S13 Histogram Coding of Nucleotide Triplets.

## Machine Learning Methods for Virus Sequence Data Analysis

### Median Neural Gas for Data Compression

The *Median Neural Gas* algorithm (MNG) is a neural data quantization algorithm for data compression based on dissimilarities [56,57]. It is a stable variant of the k-median centroid method improved by neighborhood cooperativeness enhanced learning, where *k* is the predefined number of representatives [58,59]. In this context, median approaches only assume a dissimilarity matrix for the data and restrict the data centroids to be data points. Thus, after training, MNG provides *k* data points to serve as representatives of the data. Thereby, the data space is implicitly sampled according to the underlying data density in consequence of the so-called magnification property of neural gas quantizers [60, 61].

It should be emphasized that despite the weak assumption of a given similarity matrix, MNG always delivers exact data objects as representatives. Hence, any averaging for prototype generation like in standard vector quantizers is avoided here. This is essential, if averaged data objects are meaningless like for texts, music data, or RNA/DNA sequences, for example.

### Affinity Propagation for Clustering with Cluster Number Control

*Affinity Propagation* (AP) introduced by Frey&Dueck in [62] is an iterative cluster algorithm based on message passing where the current cluster nodes, in the AP setting denoted as prototypes or exemplars, interact by exchanging real-valued messages. Contrary to methods like c-means or neural maps, where the number *c* of prototypes has to be chosen beforehand, AP starts assuming that all *N* data points are potential exemplars and reduces the number of valid prototypes (cluster centroids) iteratively. More precisely, AP realizes an exemplar-dependent probability model where the given similarities *ϛ* (*i, k*) between data points **x**_*i*_ and **x**_*k*_ (potential exemplars) are identified as log-likelihoods of the probability that the data points assume each other as a prototype. For example, the similarities *ϛ* (*i, k*) simply could be negative dissimilarities like the negative Euclidean distance.

The cost function *C_AP_* (*I*) minimized by AP is given by

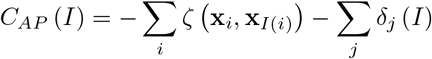

where *I*: *N* → *N* is the mapping function determining the prototypes for each data point given in (4) and

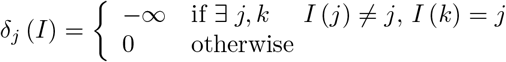

is a penalty function. During the optimization, two kind of messages are iteratively exchanged between the data until convergence: the responsibilities *r* (*i, k*) and the availabilities *a* (*i,k*). The responsibilities

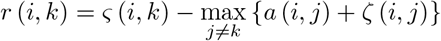

reflect the accumulated evidence that point *k* serves as prototype for data point *i*. The availabilities

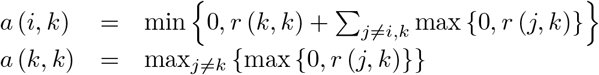

describe the accumulated evidence how appropriate data point *k* is seen as a potential prototype for the points *i*. Finally, the prototypes are determined by

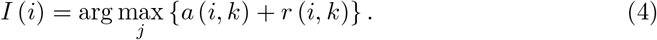

Hence, *a* (*i, k*) and *r* (*i, k*) can be taken as log-probability ratios [62]. The iterative alternating calculation of *a* (*i, k*) and *r* (*i, k*) is caused by the max-sum-algorithm applied for factor graphs [63], which can further be related to spectral clustering [64].

The number of resulting clusters is implicitly determined by the self-similarities *ϛ* (*k, k*) also denoted as preferences. The larger the self-similarities the finer is the granularity of clustering [62]. Common choices are the median or the minimum of the similarities between all inputs. Otherwise, the self-similarities can be seen as a control parameter for the granularity of the clustering. Variation of this parameter provides information regarding stable cluster solutions in dependence of plateau regions of the resulting minimum cost function value.

### Interpretable Prototype-based Classifier – the Generalized Learning Vector Quantizer

*Learning Vector Quantization* (LVQ) was is an adaptive prototype-based classifier introduced by T. Kohonen [65]. A cost-function-based variant is known as *generalized LVQ* [66]. This cost function approximates the classification error [67]. In particular, an LVQ classifier requires training data 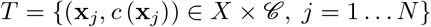 where *X* ⊆ ℝ^*n*^ and 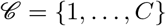 is the set of available class labels. Further, the model assumes a set of prototypes *W* = {**w**_*k*_ ∈ ℝ^*n*^, *k* = 1… *M*} with class labels *c* (**w**_*k*_) such that at least one prototype is assigned to each class. Hence, we have a partitioning of the prototype set 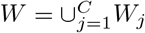 with *W_j_* = {**w**_*k*_ ∈ *W*|*c* (**w**_*k*_) = *j*}. Further, a dissimilarity measure *d* (**x, w**) is supposed, which has to be differentiable with respect to the second argument. For a given LVQ-configuration a new data point **x** is assigned to a class by the mapping

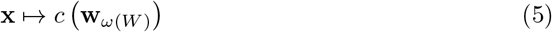

with

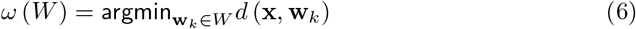

is known as the winner-takes-all rule (WTA) in prototype-based vector quantization. The prototype **w**_*ω*_ is denoted as winner of the competition.

During the learning, the cost-based LVQ minimizes the expected classification error *E_X_* [*E* (**x**_*k*_, *W*)] where

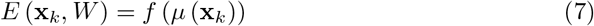

is the local classification error depending on the choice of the monotonically increasing function *f* and the classifier function

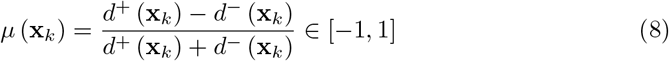

where *d*± (**x**_*k*_) = *d*± (**x**_*k*_, **w**±) and 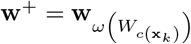 is the so-called best matching correct prototype and 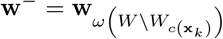 is the corresponding best matching incorrect prototype. Frequently, the squashing function *f* is chosen as sigmoid: 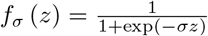. Learning takes place as stochastic gradient descent learning (SGDL) [68,69] of *E_X_* [*E* (**x**_*k*_, *W*)] with respect to the prototype set W to obtain an optimum prototype configuration in the data space.

The dissimilarity *d* (**x, w**) can be chosen arbitrarily supposing differentiability with respect to **w** to ensure SGDL. Frequently, the squared Euclidean distance *d_E_* (**x, w**) = (**x — w**)^2^ is applied resulting in the *standard generalized LVQ* (GLVQ). If both, **x** and **w** are assumed as discrete representations of density functions, divergences like the Kullback-Leibler-divergence *D_KL_* (**x, w**) from (3) come into play instead [70]. The resulting LVQ variant is denoted as *divergence-based GLVQ* (GDLVQ).

Another popular choice is the squared Euclidean mapping distance

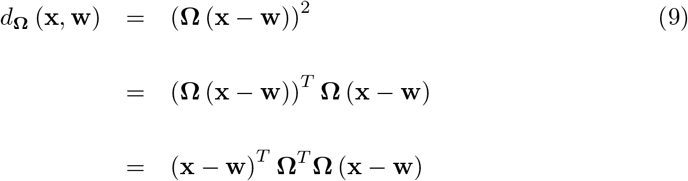

proposed in [71] with the mapping matrix **Ω** ∈ ℝ^*m×n*^ and *m* being the projection dimension usually chosen *m ≤ n* [72]. Here, the data are first mapped linearly by the mapping matrix and then the Euclidean distance is calculated in the mapping space ℝ^*m*^. The mapping matrix can be optimized again by SGDL to achieve a good separation of the classes in the mapping space. The respective algorithm is known as *Generalized Matrix LVQ* (GMLVQ) [73]. Note that SGDL for **Ω**-optimization usually requires a careful regularization technique [74].

After training, the adapted projection matrix **Ω** provides additional information. The resulting matrix **Λ** = **Ω**^*T*^**Ω** ∈ ℝ^*n×n*^ allows an interpretation as *classification correlation matrix,* i.e. the matrix entries Ω_*ij*_ give the correlations between data features *i* and *j*, which contribute to the class discrimination [39,75]. A non-linear mapping could be realized applying kernel distances instead [76, 77].

A trained LVQ model can be applied to newly incoming data of unknown distribution. However, care must be taken to ensure that the model remains applicable and that there is no inconsistency with the new data. Therefore, each LVQ can be equipped with a reject option for the application phase [78, 79]. If the dissimilarity of the best matching prototype to a data point is greater than a given threshold *τ*, it is rejected for classification, i.e. this optional tool equips the LVQ with a so-called *self-controlled evidence* (SCE) [45]. The threshold *τ* is determined during model training for each prototype individually, e.g. 95%-percentile of the dissimilarity value for those data, which are assigned to the considered prototype by the WTA-rule (6) together with the class assignment (5).

### Stochastic Neighbor Embedding for Visualization

The method of Stochastic Neighbor Embbeding (SNE) was developed to visualize high-dimensional data in a typically two-dimensional visualization space [80]. For this purpose, each data point **x**_*k*_ in the data space is associated with a visualization vector **v**_*k*_ ∈ ℝ^2^. The objective of the respective embedding algorithm is to distribute the visualization data in a way that the density of original data distances in the high-dimensional data space is preserved as good as possible for the respective density of the distances in the visualization space (embedding space). The quality criterion is the Kullback-Leibler-divergence between them, which is minimized by SGDL with respect to the visualization vectors **v**_*k*_.

Yet, SNE suffers from the fact that the distance densities in the original data space are frequently heavy-tailed [81], which leads to inaccurate visualizations. To overcome this problem, the so-called *t-distributed* SNE (t-SNE) was developed [82].

### Data Processing Workflow

In the following we describe and motivate the steps of data processing and analysis.

1. Coding of all sequences of *D_G_*-data and *D_N_*-data.

- Alphabet 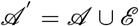 with alphabet extension 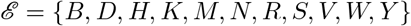 due to ambiguous characters in the data sets.
- A natural vector representation **x** (4, **s**) ∈ ℝ^20^ of order *K* = 4 is generated for each sequence **s** according to (1) paying attention to the alphabet extension 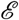.
- A BoW-representation for 3-mers is generated for each sequence **s**: **h** (**s**) ∈ ℝ^64^ according to the possible nucleotide triplets of the alphabet 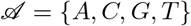 paying attention to the alphabet extension 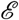
2. Training of LVQ-classifiers for *D_G_*-data to evaluate the results from [12] obtained by phylogentic trees

- Training data are all samples of *D_G_* with the additional virus type assignment *A, B*, or *C* taken as class labels.
- For all LVQ variants we take only one prototype per class.
- For GMLVQ, the projection matrix is chosen as **Ω** ∈ ℝ^2×*n*^, i.e. the mapping dimension is *m* = 2.
- SGDL training as 10-fold cross validation to determine the best LVQ architecture for the given problem.

— Training of *W* using the GLVQ for NV representation. * GDLVQ is not applicable for this sequence representation due to mathematical reasons.
— Training of *W* using the GLVQ for BoW representation.
— Training of *W* and **Ω** using the GMLVQ for BoW representation.
- Final training of the best LVQ architecture with optimum training schedule to achieve best prototype configuration *W*.

— If GMLVQ architecture is selected for final training: training of both W and **Ω**, determination of the classification correlation matrix **Λ** = **Ω**^*T*^**Ω**.
— Determination of the reject thresholds for each prototype for self-controlled of evidence use based on the 95%-percentile rule.
3. Clustering *D_N_*-data

- Compression of the subset of 706 US-sequences of March by MNG to achieve 50 representatives by MNG using 50 prototypes.
- generating a balanced subsets consisting of all China samples (63), all Europe samples (33), and USA samples (114) for cluster analysis. The US samples comprise the 50 representatives from MNG and all US samples from January and February. The sample from other regions are not considered for cluster analysis. We denote this balanced data set extracted from *D_N_* by *D_NB_*.
- Clustering and identification of stable cluster solutions using affinity propagation by means of the control parameter *ϛ* = *ϛ* (*k, k*) ∀*k*.
4. Classification of the *D_NB_*-data as well as the full *D_N_*-data using the best LVQ classifier with integrated self-controlled evidence

- Classification of the *D_NB_* -data by the final LVQ classifier with reject option using the determined thresholds to realize the self-controlled evidence (SCE).
- Evaluation of the data rejected by the SCE rule.

## Results

According to the processing workflow we trained several LVQ-classifier variants for the *D_G_*-data. By 10-fold cross-validation, we achieved the averaged accuracies depicted in Tab. 2 together with their respective standard deviations. According to these results, GMLVQ performs best using the BoW-coding of the sequences together with the Euclidean mapping distance *d*_Ω_ (**x, w**) from (9). Thus, we finally trained a GMLVQ network for both the prototype set W containing one prototype per class and the mapping matrix **Ω** using the sequence BoW-coding. For this final network a classification accuracy of 100% is obtained while rejecting 7 samples for classification according to the SCE-decision. The resulting classification correlation matrix **Λ = Ω**^*T*^**Ω** is depicted in S1 Fig. Because **Ω** ∈ ℝ^2×*n*^, it can serve for a data mapping into a two-dimensional visualization space. Accordingly, all *D_G_*-data together with the GMLVQ-prototypes are visualized in S2 Fig. An additional visualization of the learned prototypes is given in S3 Fig.

The list of rejected sequences is provided in the supplementary material S14 GMLVQ Mapping for *D_N_*.

The clustering of the *D_NB_*-data set suggests cluster solutions with either 2, 4, or 5 clusters according to the stability range of the control parameter *ϛ*, see S4 Fig. We visualized the 4-cluster solution using the *t*-SNE as depicted in S5 Fig. The respective cluster centroids are visualized in S6 Fig.

**Table 2.**
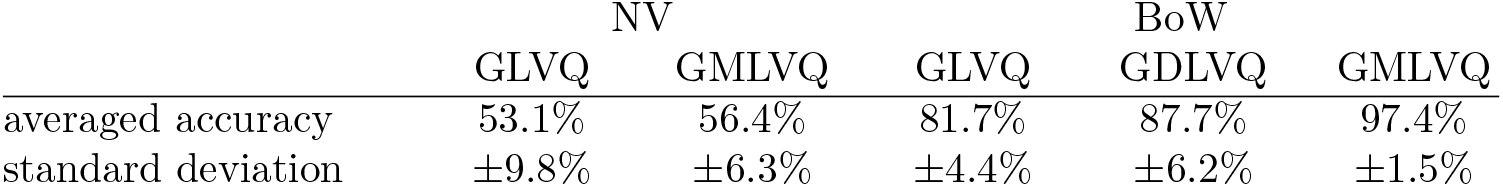
Classification results of trained LVQ-variants for the *D_G_*-dataset obtained by 10-fold cross-validation.

Applying the trained GMLVQ classifier to the *D_NB_*-data set leads to the classification of 37 data points to class *A*, 95 data points to class *B*, 2 data points to class *C*. According to the SCE-decision, 59 data points were rejected from classification by the learned GMLVQ classifier. The result is given in S7 Fig using the *t*-SNE as visualization scheme. The visualization of the classification result by means of the **Ω**-mapping from the GMLVQ model delivers S8 Fig.

The distribution of the sequence data from the *D_NB_*-data set with respect to the geographic sequence origins (regions) and the respective collection dates together with the class assignments is presented in S9 Fig. A respective visualization of the distribution for the data set *D_G_* is shown in S10 Fig.

The classification of the full *D_N_* data set assigns 154 data points to class A, 293 data points to class *B*, and 20 data points to class *C*, whereas 495 data points are rejected according to the SCE-rule. The class assignments are visualized in S11 Fig.

The predicted virus type or the rejection decision for each sequence from *D_N_* according to the GMLVQ class assignment or the RCE decision can be found in the supplementary material S14 GMLVQ Mapping for *D_N_*.

## Discussion

The classification analysis of the *D_G_*-data by means of the machine learning model GMLVQ verifies the class determination suggested in [12]. Only 7 data samples are not classified accordingly due to the model self-controlled evidence decision. Thereby, the GMLVQ model shows a stable performance in learning (see Tab. 2), which underlies its well-known robustness [47]. Thus, we observe an overall precise agreement supporting the findings in [12].

This agreement, however, is obtained by alignment-free sequence comparisons. More precisely, the nucleotide based BoW sequence coding delivers a perfect separation of the given classes for the learned mapping distance *d*_Ω_ (**x, w**). Yet, the computational complexity of dissimilarity calculations for the encoded sequences is only *O* (64 · *m*) with *m* = 2 being the mapping dimension of **Ω**. The BoW sequence coding takes *O* (*n*) such that we have an overall complexity of *O* (*n* + 64 · *m*) = *O* (*n*) for dissimilarity calculations needed in our approach in comparison to at least *O* (*n*)^2^ in case of alignment based dissimilarity calculations.

Because GMLVQ is an interpretable classifier, we can draw further conclusions from the trained model: The resulted classification-correlation matrix **Λ** depicted in S1 Fig suggests that particularly the histogram dimensions 27 and 28 are important in correlation with the other dimensions. These dimensions refer to the frequency of the triplets ‘CGG’ and ‘CGT’ in the sequences. Moreover, both dimensions should be negatively correlated for good class separation. This discrimination is a key feature of GMLVQ. Although the prototypes look very similar, see S3 Fig, the **Ω** is sensitive tosmallest deviations in the histograms. Yet, we cannot expect greater deviations, because the sequences differ only in few characters according to the special mutations [11, 12]. The AP-centroids differ slightly more than the GMLVQ prototypes, see S6 Fig. This can be dedicated to larger overall scattering of the *D_NB_*-data.

Further, the GMLVQ-prototypes serve as class ‘detectors’. If the encoded sequences are most similar to them with respect to the mapping distance, the sequences are assigned to the respective classes according to the WTA (6). However, in general the prototypes are not identical with the mean vectors of the class distribution, as emphasized in [83].

Application of the GMLVQ to the *D_N_*- and *D_NB_*-data from the NCBI offers new insights. First, coloring of the data in the t-SNE visualization S7 Fig of *D_NB_* according to the obtained class assignments seems to be confusing: The classes can not be detected as separate regions in that case. However, applying the **Ω**-mapping S8 Fig, the class structure becomes visible also for this data set. The reason for this discrepancy could be that both *t*-SNE as well as AP implicitly reflect data densities in the data space. Class densities, however, do not have to coincide with the overall data density.

Thus, the **Ω**-mapping, which is optimized during GMLVQ training for best classification performance, offers the better visualization option and, hence, disclosures the class distribution more appropriately.

Comparing the class distributions of the sequences with respect to origins (regions) and collection dates for *D_NB_* in S9 Fig and *D_G_* in S10 Fig both class distributions within the cells show a similar behavior. The *D_NB_*-data set from NCBI contains only a few samples from Europe all occurring from February onward, i.e. no European data samples from December/January are available. We observe that class *C* for the *DG* data is mainly represented in January for European samples, which confirms the findings in [12]. Thus, the small number of class *C* samples in *D_NB_*-classification may be addressed to this peculiarity in Europe. Further, the GMLVQ, which was trained by *D_G_* data, rejects a large amount of data from *D_NB_*, particularly in March. We suspect an accumulation of mutations which could explain the scattering. Accordingly, the GMLVQ is able to detect this behavior by means of the SCE decision rule.

We observe from the visualization S11 Fig of the classification for the *D_N_*-data that the data points rejected for classification scatter around the dense class regions. Thus we can conclude that the nucleotide base mutations in the viral sequences, which cause the scattering, do not show a new coherent profile, at least at this time.

## Conclusion

In this contribution we investigate the application of interpretable machine learning methods to identify types of SARS-CoV-2 virus sequences based on alignment-free methods for RNA sequence comparison. In particular, we trained a *Generalized Matrix Learning Vector Quantizer* (GMLVQ) classifier model (GMLVQ) for a data set with given virus type information, which was obtained by phylogenetic tree analysis [12]. GMLVQ supposes vectorial data representations and compares vectors in terms of a well-defined dissimilarity measure. In this application, the GMLVQ training is based on the Bag-of-Words coded sequences and yields class specific prototype vectors as well as an optimum class/typus separating dissimilarity measure in the data space of encoded sequences. Compared to phylogentic trees, which require high computational costs due to the involved sequence alignment process, the GMLVQ approach has lower complexity and allows an easy out-of-training generalization.

By means of the trained GMLVQ we first verified the SARS-CoV-2 virus types determined in this first data set. Further, considering a classification correlation matrix delivered by GMLVQ optimization, we are able to identify features which contribute decisively to a type separation.

Second, we applied the trained GMLVQ to another data set obtained from the NCBI database without virus type information. Using the self-controlled evidence property of the GMLVQ we are able to classify these sequences to the previously identified types avoiding the application of the model to inconsistent data compared to the training data. Further, the rejected data allow speculations about new virus types with respect to nucleotide base mutations in the viral sequences.

Future work will consider to replace the WTA-rule (6) by a fuzzy variant (winner ranking) resulting in a probabilistic class/type assignment replacing the crisp rule (5). This probabilistic view could be further integrated into the SCE based rejection decision to differentiate between rejected sequences regarding their consistence to the GMLVQ version in use. Thus, the user can decide whether to retrain the model adding a new class or continue with the current configuration.

## Supporting information

**S1 Fig.**
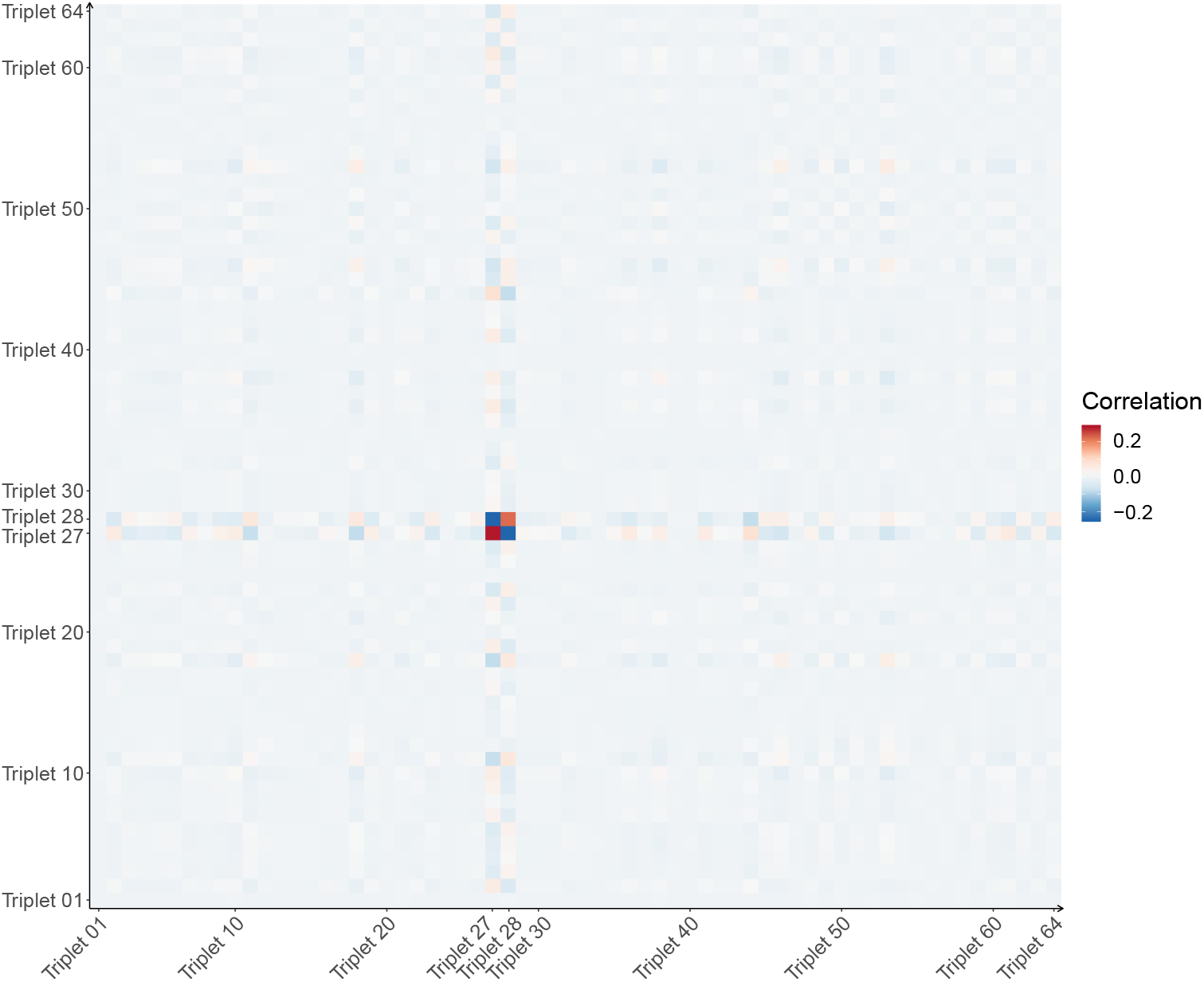
Visualization of the classification-correlation matrix. The classification-correlation matrix provides the linear correlations among the triplet frequencies of the sequences, which contribute to the class discrimination.

**S2 Fig.**
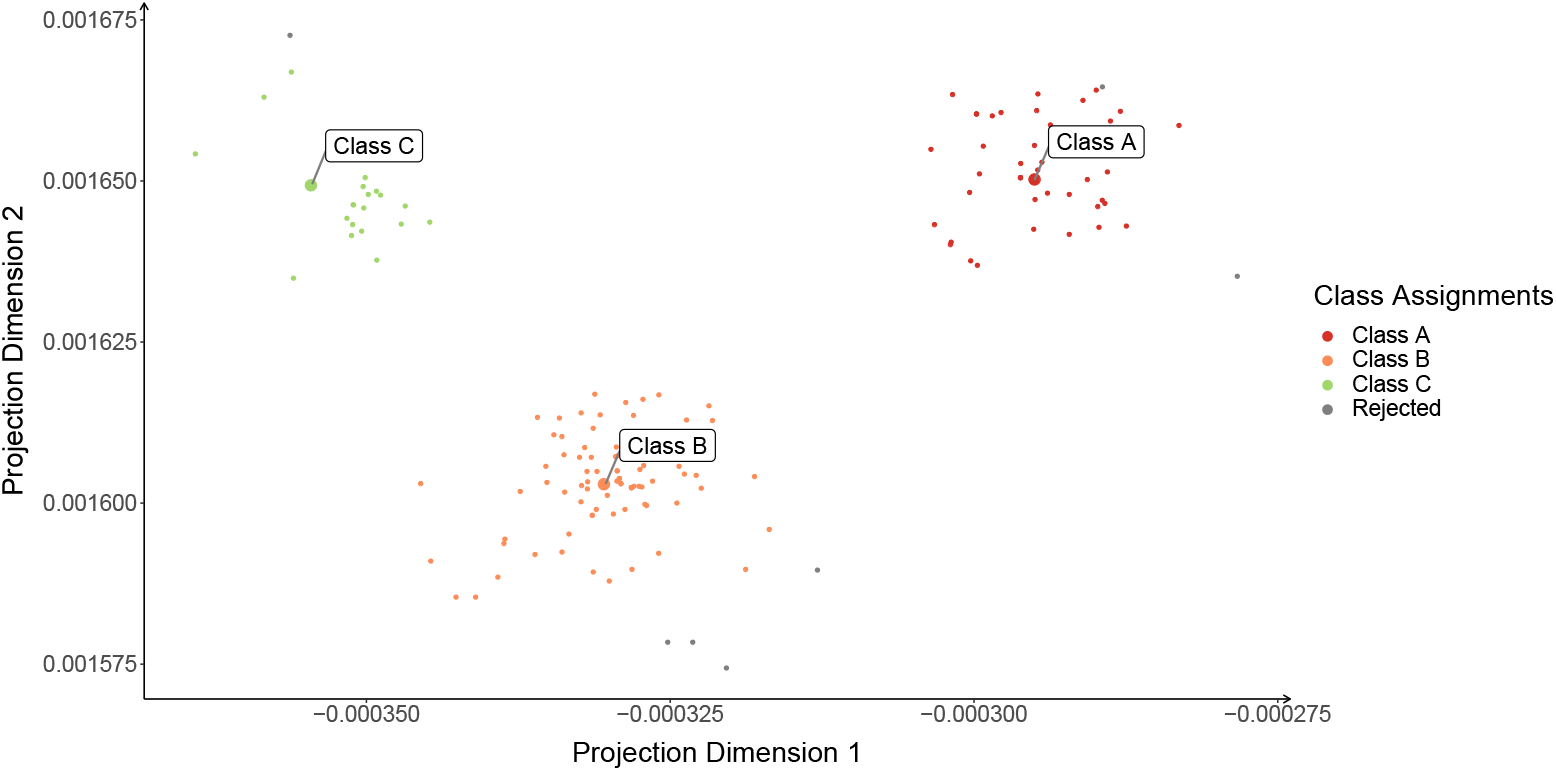
Visualization of the GMLVQ result for *D_G_*-data. The data as well as the the GMLVQ-prototypes are mapped using the learned **Ω**-matrix. The data points are colored either regarding their class assignments or regarding their reject decision. The GMLVQ-prototypes serve as class representatives. However, they are not identical with the mean vectors of the classes.

**S3 Fig.**
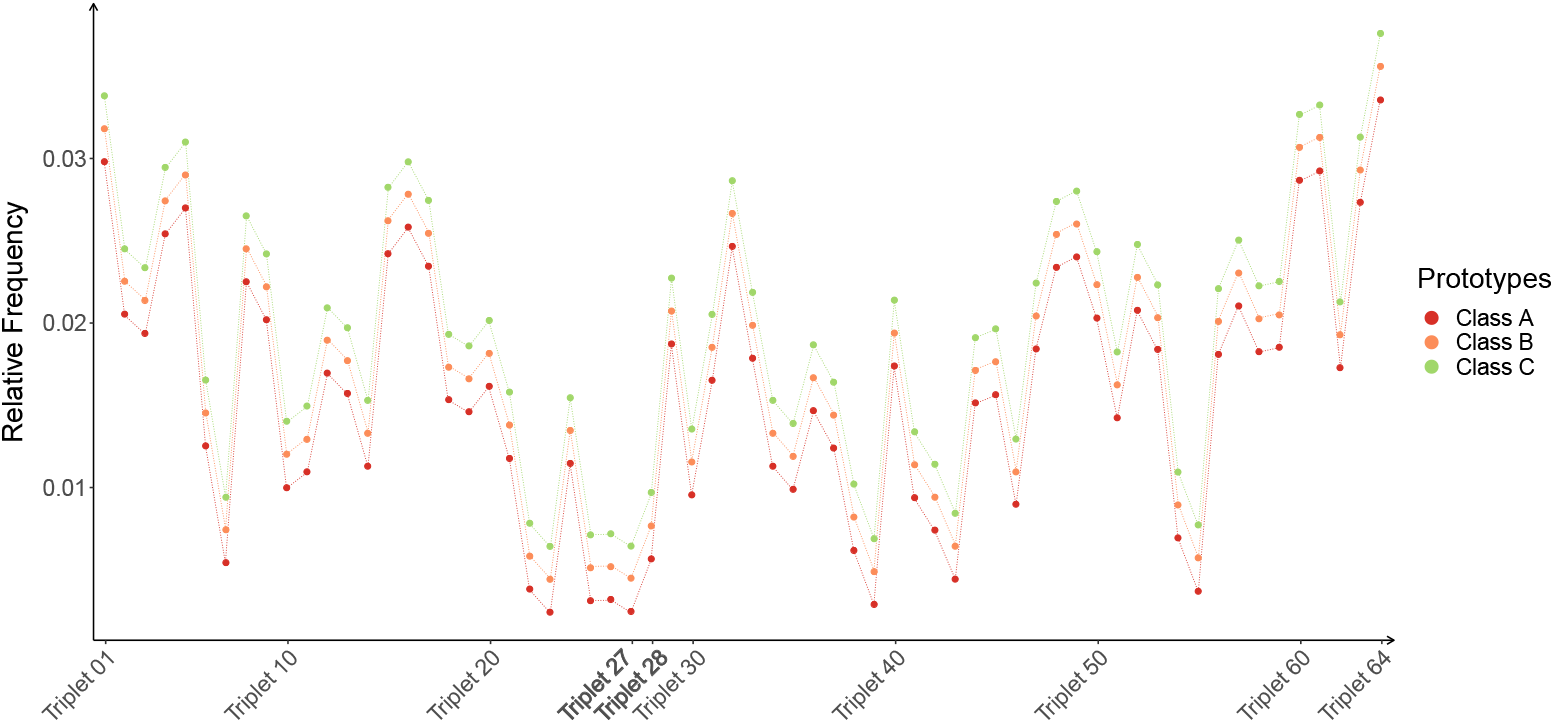
Visualization of the GMLVQ prototypes. The color of the prototypes is in agreement with the class coloring in S2 Fig. Further, the prototypes are vertically shifted by small offsets for better visualization and separation.

**S4 Fig.**
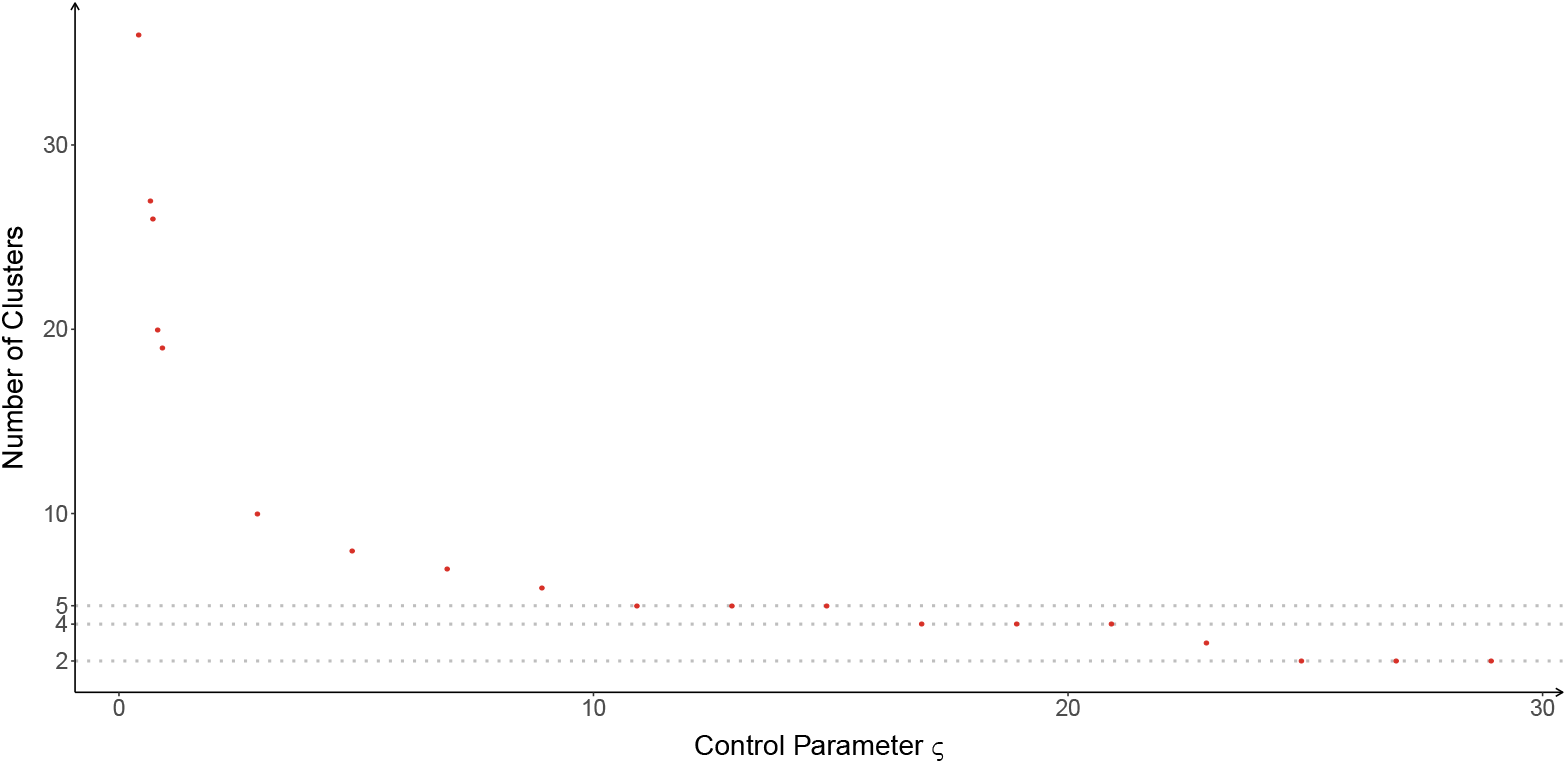
Stability of AP cluster solutions for *D_NB_*-data. The number of clusters in dependence on the control parameter *ϛ* is depicted. Plateaus refer to stable cluster solutions. Accordingly, we identify 2-, 4-, and 5-cluster solutions as most recommended.

**S4 Fig.**
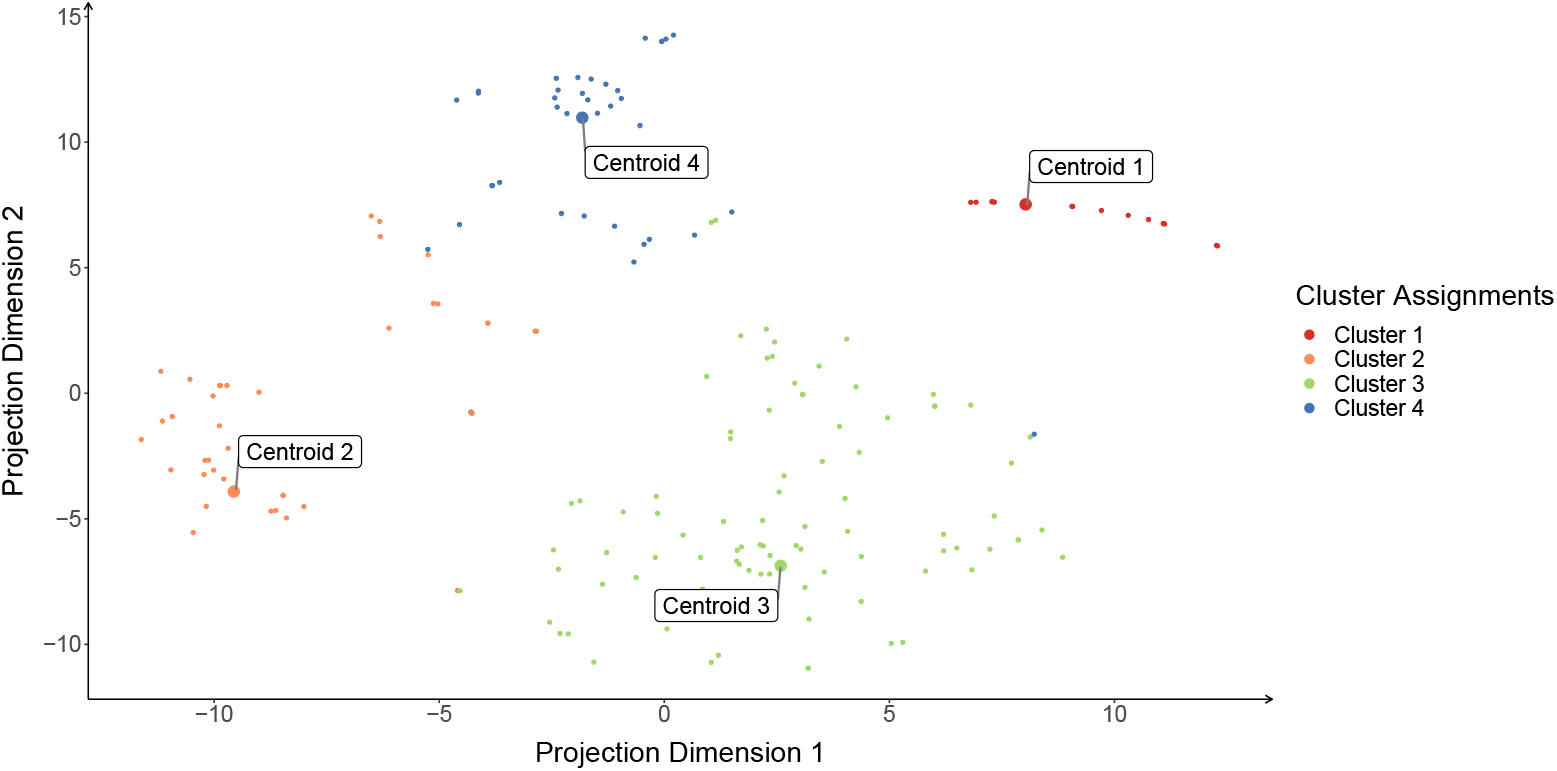
Visualization of the AP clustering result for *D_NB_*-data using 4 clusters. The data as well as the cluster centroids are depicted using the *t*-SNE.

**S6 Fig.**
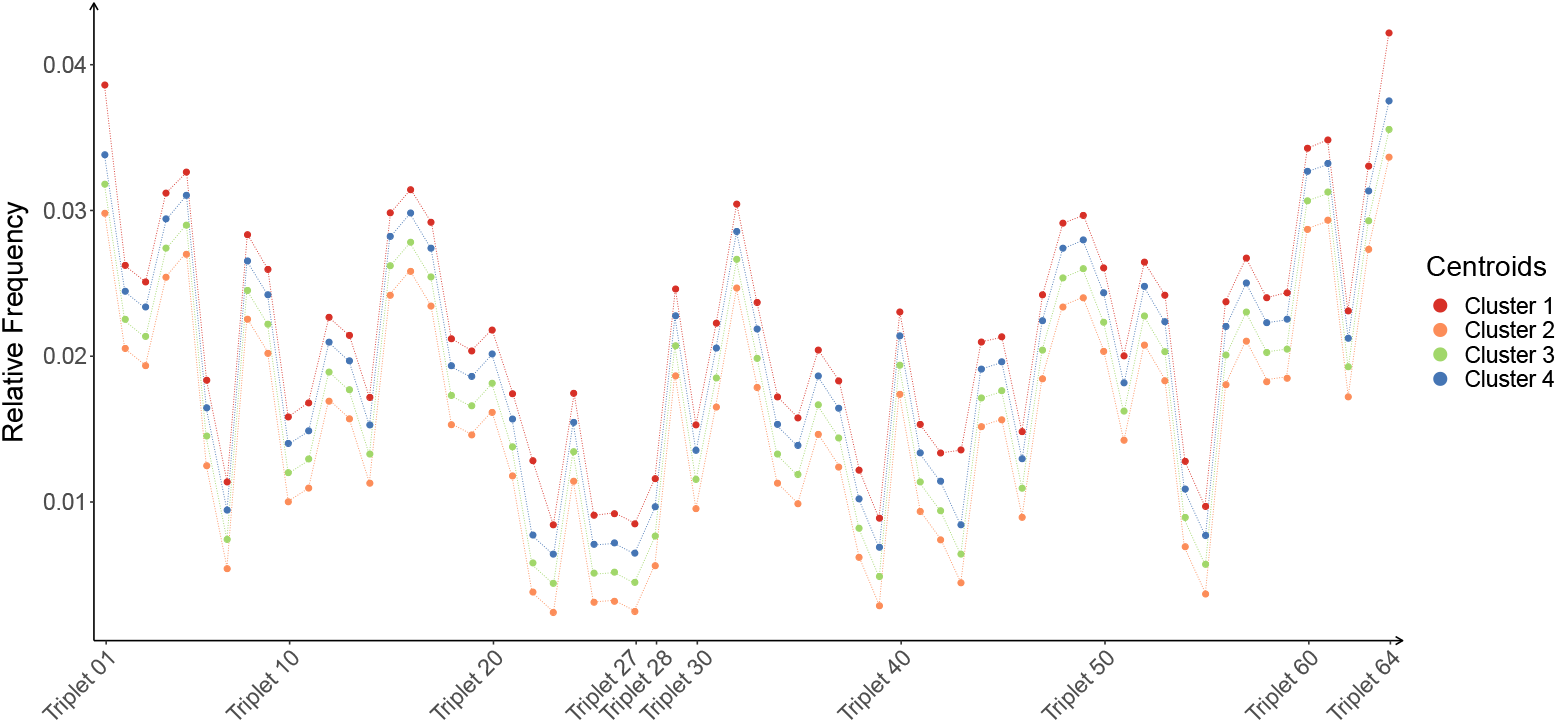
Visualization of the AP cluster centroids for *D_NB_*-data using the 4-cluster solution. The color of the cluster centroids is in agreement with the cluster coloring in S5 Fig. Further, the centroids are vertically shifted by small offsets for better visualization and separation.

**S7 Fig.**
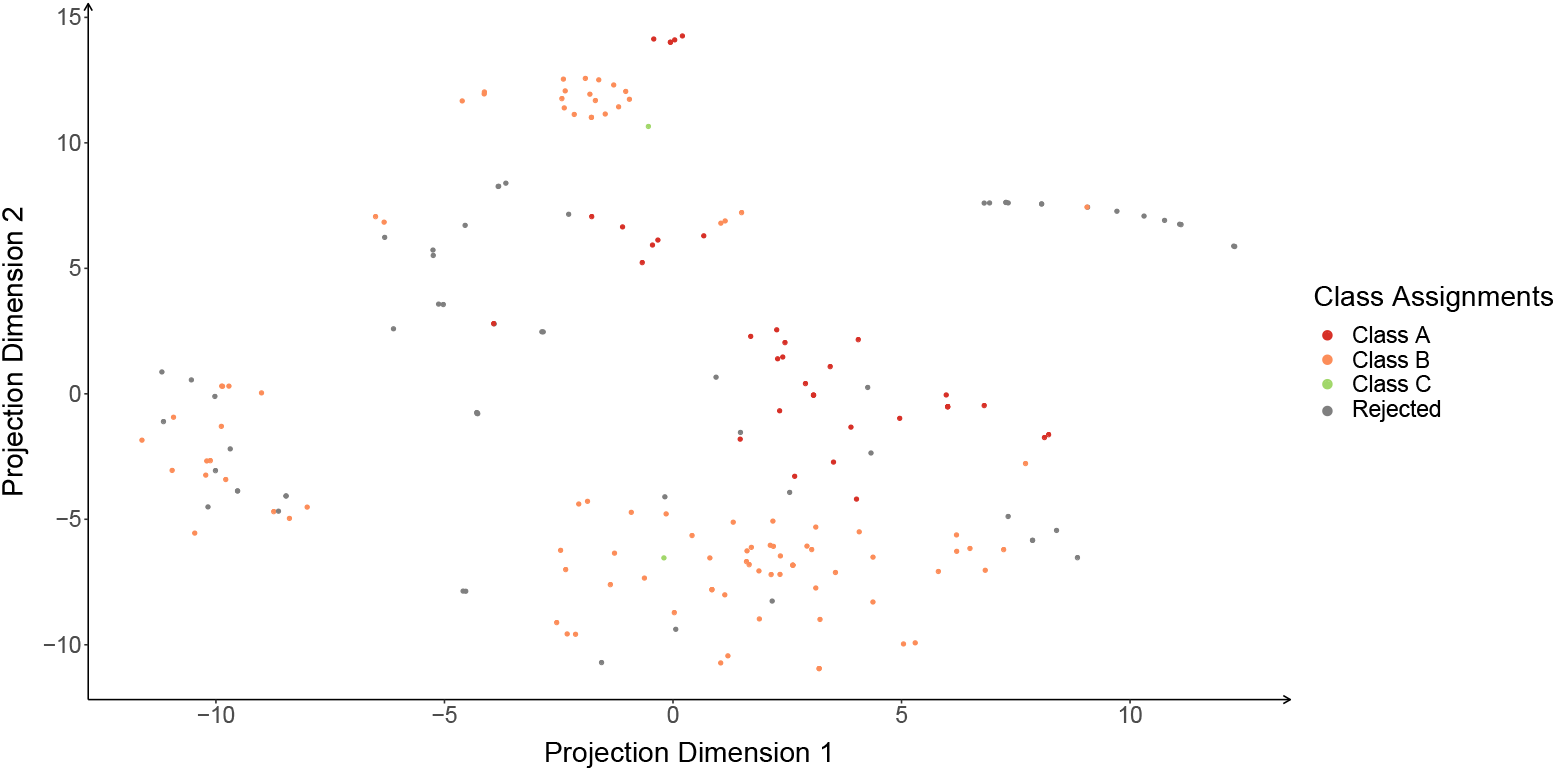
Visualization of GMLVQ classification for the *D_NB_*-data by t-SNE. The class coloring is as in S2 Fig.

**S8 Fig.**
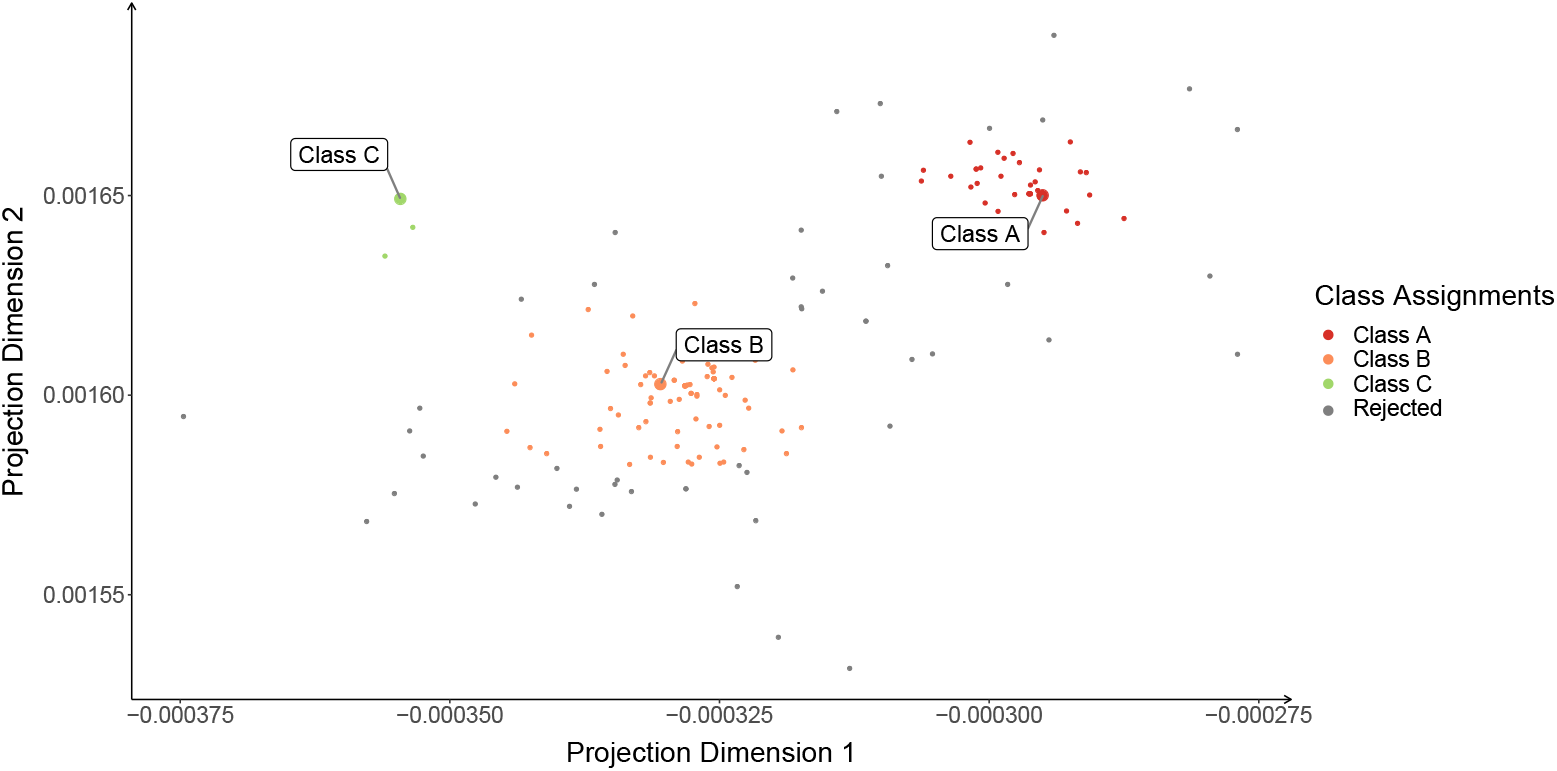
Visualization of GMLVQ classification for the *D_NB_*-data by Ω-mapping. The data as well as the GMLVQ prototypes are depicted. The class coloring is as in S2 Fig.

**S9 Fig.**
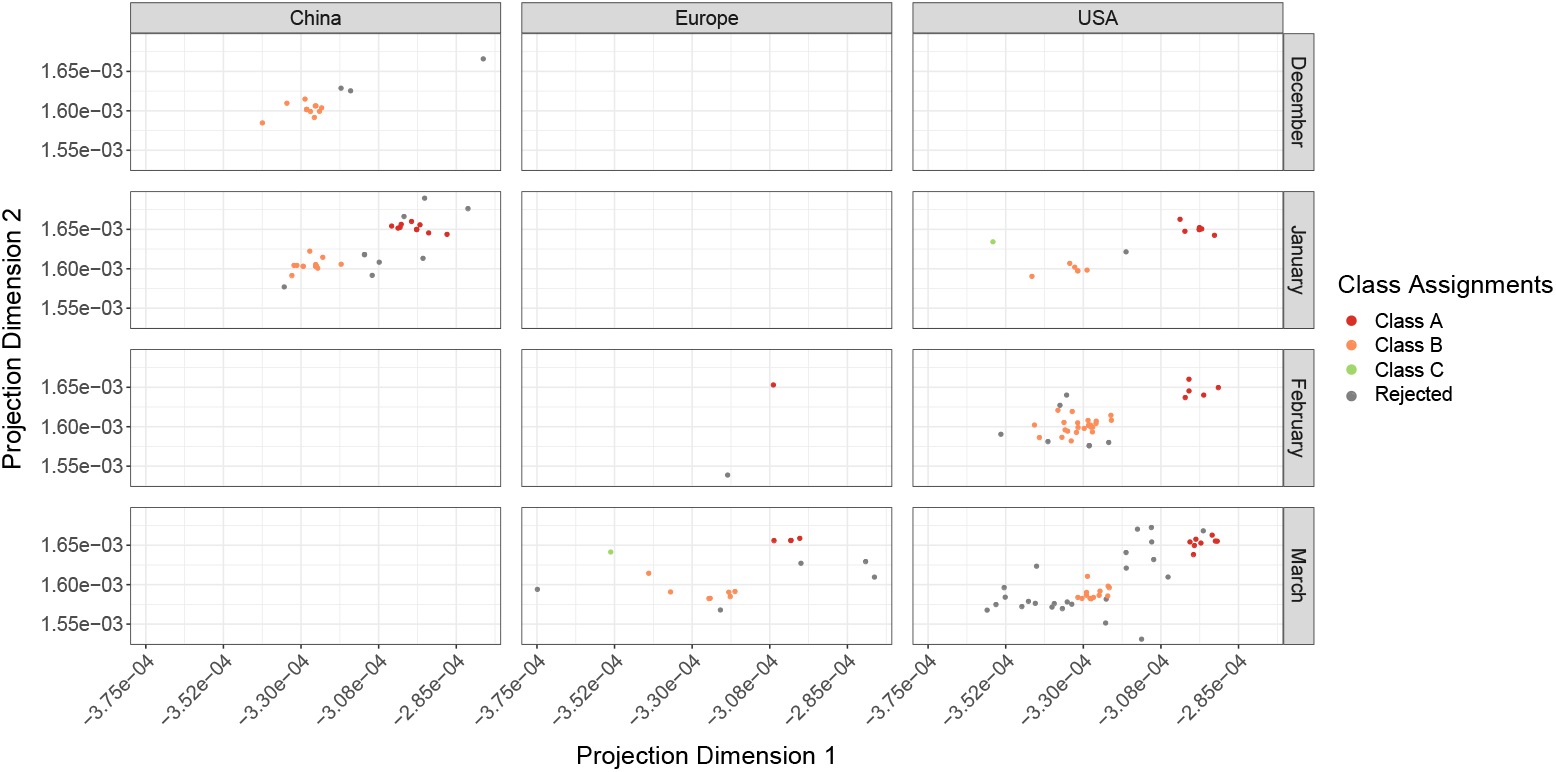
Distribution of the *D_NB_*-data depending on the geographic origin and the collection date. A distribution of the data from the *D_NB_*-data set with respect to the geographic sequence origin and the collection date together with the class assignments. Again, the mappings are realized by the **Ω** matrix. The class coloring is as in S2 Fig.

**S10 Fig.**
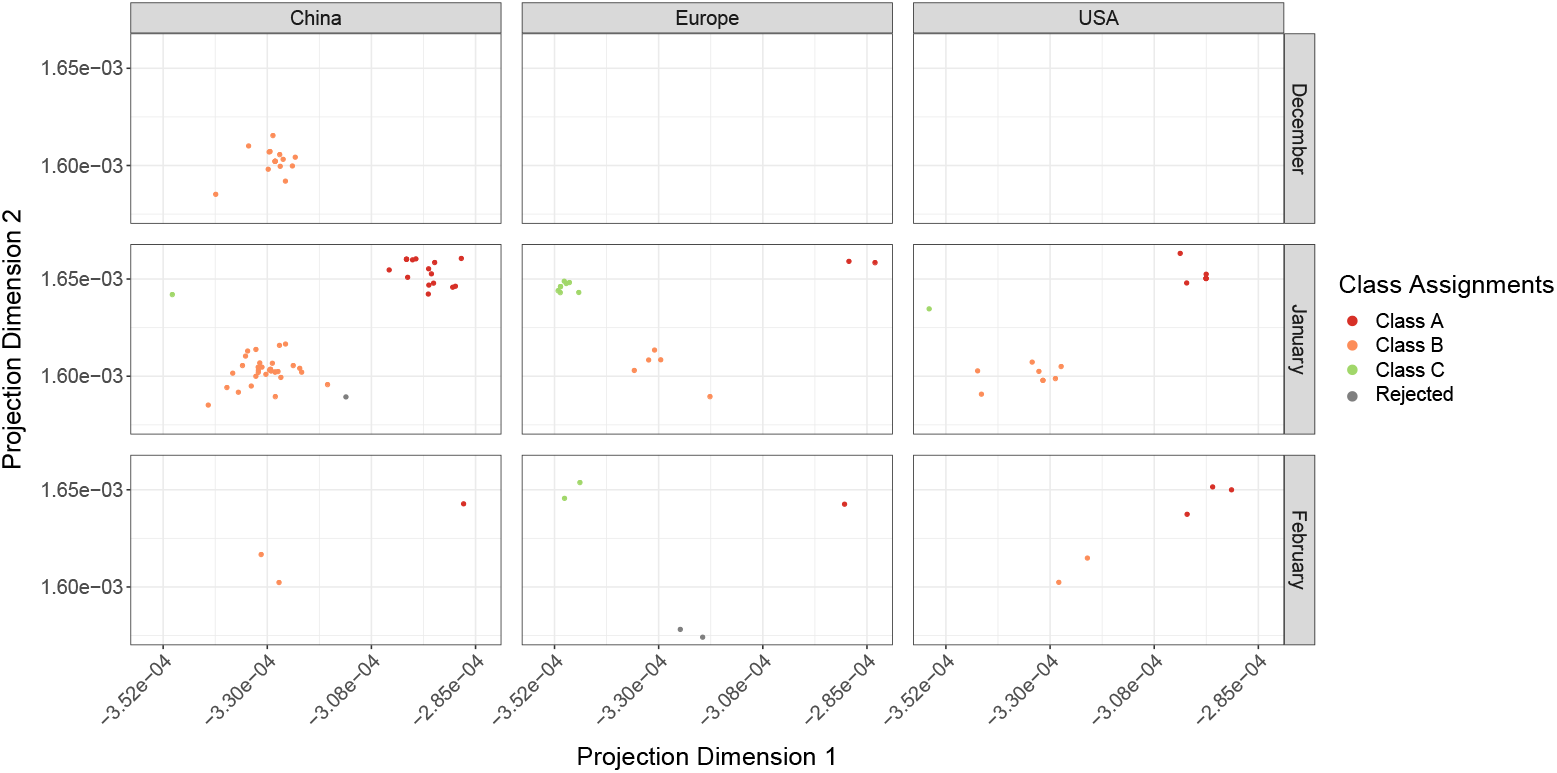
Distribution of the *D_G_*-data depending on the geographic origin and the collection date. A distribution of the data from the *D_G_*-data set with respect to the geographic sequence origin and the collection date together with the class assignments. Again, the mappings are realized by the **Ω** matrix. The class coloring is as in S2 Fig.

**S11 Fig.**
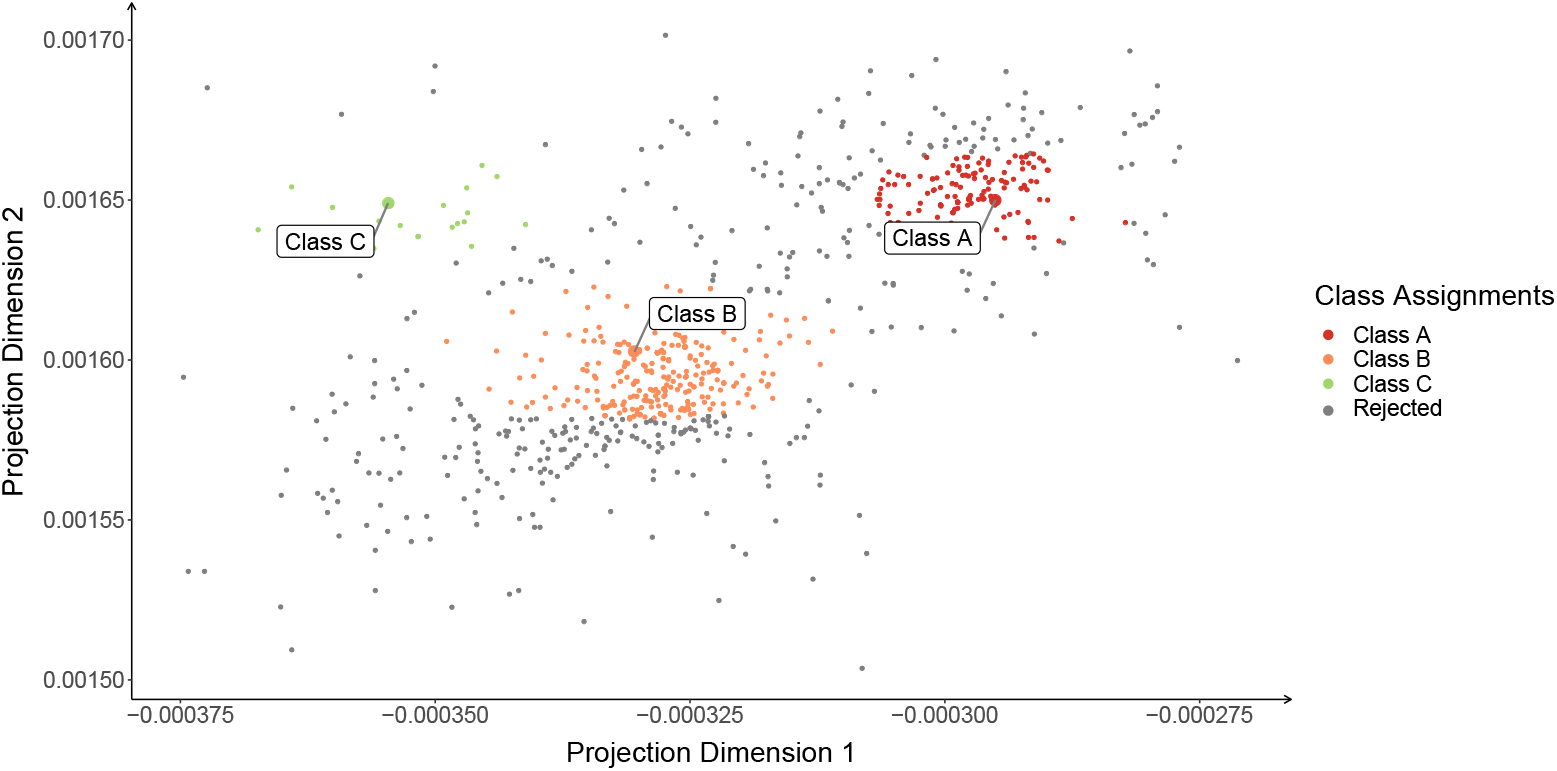
Visualization of GMLVQ classification for the full *D_N_*-data by Ω-mapping. The data as well as the GMLVQ prototypes are depicted. The class coloring is as in S2 Fig.

**S12 Data Data Files** The data file ‘S12 Data.xlsx’ (Excel) contains the accession numbers of both data sets *D_G_* and *D_N_*. Further collection date and origin (region) are attached. For the DG data set, additionally, the type information (class assignment) is given.

**S13 Histogram Coding of Nucleotide Triplets Assignment of the histogram dimensions to the nucleotide triplets** For each of the 64 nucleotidecombinations, the coding by the histogram dimensions is given in the file ‘S13 Histogram Coding.xlsx’.

**S14 GMLVQ Mapping for** *D_N_* **Virus type assignments for the** *D_N_*-**sequences obtained by GMLVQ** For each sequence in *D_N_*, the class/type assignment obtained by the GMLVQ model is given as well as if a rejection decision was made according to the SCE of the GMLVQ model. Additionally, we provide the sequences from *D_G_*, which were rejected by the SCE decision of the GMLVQ model. The respective file is ‘S14 GMLVQMapping.xlsx’.

## Acknowledgments

M.K., K.S. B., M.W., and M.K. acknowledge support by a grant of the European Social Fund (ESF).

